# The Winner Takes It All: a single genotype of *Kalanchoe* ×*houghtonii* is a global invader

**DOI:** 10.1101/2025.02.03.636197

**Authors:** Joan Pere Pascual-Díaz, Neus Besolí, Jordi López-Pujol, Neus Nualart, Iván Pérez-Lorenzo, Ronen Shtein, Laura Valenzuela, Sònia Garcia, Daniel Vitales

## Abstract

**Background and Aims:** Invasive alien plant species pose a global challenge, and their impact is amplified by globalisation and the accelerating pace of climate change. In mild-climate regions, drought-tolerant invasive plants showing broad environmental tolerance have a competitive advantage. One example is *Kalanchoe* ×*houghtonii* (Crassulaceae), popularly known as “mother of millions”. It is a hybrid resulting from the interploid cross between *K. daigremontiana* and *K. delagoensis*, both native to Madagascar. *Kalanchoe* ×*houghtonii*, propagated as an ornamental plant, has emerged as a global invader in less than a century. Four morphotypes of this hybrid have been identified, with different ploidy levels and varying invasive capacities. Here we aim to investigate the genomic variability behind the invasion success of *Kalanchoe* ×*houghtonii*.

**Methods:** We sampled 57 accessions of *Kalanchoe* ×*houghtonii, K. daigremontiana, K. delagoensis* and closely related taxa, including old herbarium materials, from all over the world. We analysed genome size, chromosome numbers, sequenced the whole genome, analysed the complete plastome sequence of each accession, and studied the diversity of the ribosomal RNA genes. We also performed a detailed phylogenomic study using nuclear BUSCO genes.

**Key Results:** Our study reveals the genetic and cytogenetic variability between morphotypes, and shows that a single tetraploid genotype (morphotype A) dominates all populations, emerging as the first reported clonal hybrid capable of worldwide colonisation. Morphotype A shows a striking genetic uniformity, high phenotypic plasticity, and extremely high rates of vegetative reproduction, representing an example of a “general-purpose genotype”.

**Conclusions:** The astonishing reproductive capacity, broad adaptability and the speed at which *K.* ×*houghtonii* is colonising new regions by clonal spread highlight the importance of understanding hybridisation and polyploidy in the invasion of ecosystems. Our findings call for the need for risk assessments before developing new hybrids for ornamental plant breeding that may exhibit invasive characteristics.

## Introduction

Invasive alien plant species (IAPS) are a global problem due to their multiple environmental and economic impacts (Hulme, 2003; Bresch *et al*., 2013), and are considered a major driver of recorded species extinctions (Roy *et al*., 2023). Their expansion beyond their native range is caused by human activities and is often determined by socioeconomic drivers (Pysek *et al*., 2020), which promote their appearance even in protected or highly biodiverse ecosystems (Gioria *et al*., 2023). Although some IAPS have been accidentally introduced, most introductions are intentional, in many cases linked to the globalised trade of ornamental plants (Hulme *et al*., 2008; Pysek *et al*., 2011a; van Kleunen *et al*., 2018). Their subsequent spread and establishment can threaten ecosystems, while causing economic impacts related to the cost of their management (Binggeli and McNeely, 2001; Pysek *et al*., 2009).

The Mediterranean basin is recognized as a biodiversity hotspot (Mittermeier *et al*., 2011). The intense human activity and land degradation in this area (Hill *et al*., 2008; Underwood *et al*., 2009), together with climate change (MedECC, 2020; Herrando-Moraira *et al*., 2022; Urdiales-Flores *et al*., 2023), make the Mediterranean basin one of the most vulnerable regions to be affected by IAPS (Gritti *et al*., 2006; Cao Pinna *et al*., 2021). According to Keller *et al*. (2011), at least 1,780 non-native species have been established in natural ecosystems across Europe, including IAPS with high ecological and economic impact, such as *Ailanthus altissima*, *Cortaderia selloana* or *Robinia pseudoacacia* (Pysek *et al*., 2009) – most of them coming from the ornamental plant trade. Specifically, the Mediterranean region is harbouring at least 501 invasive taxa (Pysek *et al*., 2011*b*), however this may be already underestimated, as a more recent study reports 250 IAPS only in Italy (Galasso *et al*., 2024). As the Mediterranean is becoming warmer and drier, representatives of drought-tolerant groups of plants such as Aizoaceae, Cactaceae and Crassulaceae, typically succulent, are being increasingly established, e.g. several species of the genus *Kalanchoe* Adanson (Stinca *et al*., 2021; Sakhraoui *et al*., 2023).

*Kalanchoe* comprises approximately 140 species mainly native to Madagascar (Milad *et al*., 2013; Kuligowska *et al*., 2015). Many species of the genus are well known for their commercial value, mostly because of their ornamental and medicinal uses (Kołodziejczyk-Czepas and Stochmal, 2017; Stefanowicz-Hajduk *et al*., 2020; Vargas *et al*., 2022). Although natural hybrids are not common within the genus (Smith *et al*., 2019; Smith & Figueiredo, 2019; Smith, 2021), the ease with which hybrids can be created in cultivation has given rise to the description of several ornamental cultivars (Smith, 2020*a-g*; Smith and Figueiredo, 2020; Smith and Shtein, 2020).

*Kalanchoe* ×*houghtonii* is an alleged artificial hybrid complex resulting from the cross between the diploid species *K. daigremontiana* (2*n* = 2*x* = 34) and the tetraploid species *K. delagoensis* (2*n* = 4*x* = 68), both of native to Madagascar and growing sympatrically in some places of the island. *Kalanchoe* ×*houghtonii* has been synthetically created at least twice, once in the US by A. D. Houghton (1935) and again in Portugal by F. Resende (1954). Morphologically, *Kalanchoe* ×*houghtonii* is a perennial erect succulent herb, generally monocarpic (dying after flowering), that may reach up to 1.8 m. It has long, narrow, and triangular-shaped leaves, which are serrate and mottled (Fig. 1). It flowers annually in late winter, with inflorescences that bear over 100 pendulous dark-red flowers (Herrando-Moraira *et al*., 2020). Shtein *et al*. (2021) reported the existence of four different hybrid morphotypes (named A, B, C and D). Plants of *K.* ×*houghtonii* morphotype A are very invasive, showing an extraordinary adaptation capacity to colonise and be naturalised in numerous mild-climate regions around the world (Herrando-Moraira *et al*., 2020). Although it has been speculated that this morphotype would have been artificially produced more than once on different continents, and that it could correspond to a fertile, likely tetraploid, plant as the one Resende (1954) created in Portugal, the true origins and cytogenetic features of morphotype A are not yet known. Plants of morphotype B are significantly less invasive than those of morphotype A, though individuals representative of morphotype B have occasionally become naturalised in some places (e.g. in Spain, Shtein *et al*., 2021). The original hybrid material created by Houghton (1935) likely corresponds to morphotype B, being later cytogenetically characterised by Baldwin (1949) as a triploid. Morphotype C is suspected to be a spontaneous hybrid between *K. daigremontiana* and *K. delagoensis* in Madagascar or a closely related non-hybrid taxon showing intermediate characters (Shtein *et al*., 2021). Finally, morphotype D occurs naturally in Madagascar, where it is thought to result from the natural introgression between *K. daigremontiana* and native *K.* ×*houghtonii* plants but can also be found as a cultivar (‘Parsel Tongue’), likely stemming from a back-cross of *K. daigremontiana* with the fertile cultivar *K.* ×*houghtonii* ‘J.T. Baldwin’ (morphotype A) (Shtein *et al*., 2021). As with morphotype A, the chromosome numbers and ploidy levels of morphotypes C and D have never been determined with certainty.

**FIGURE 1.**
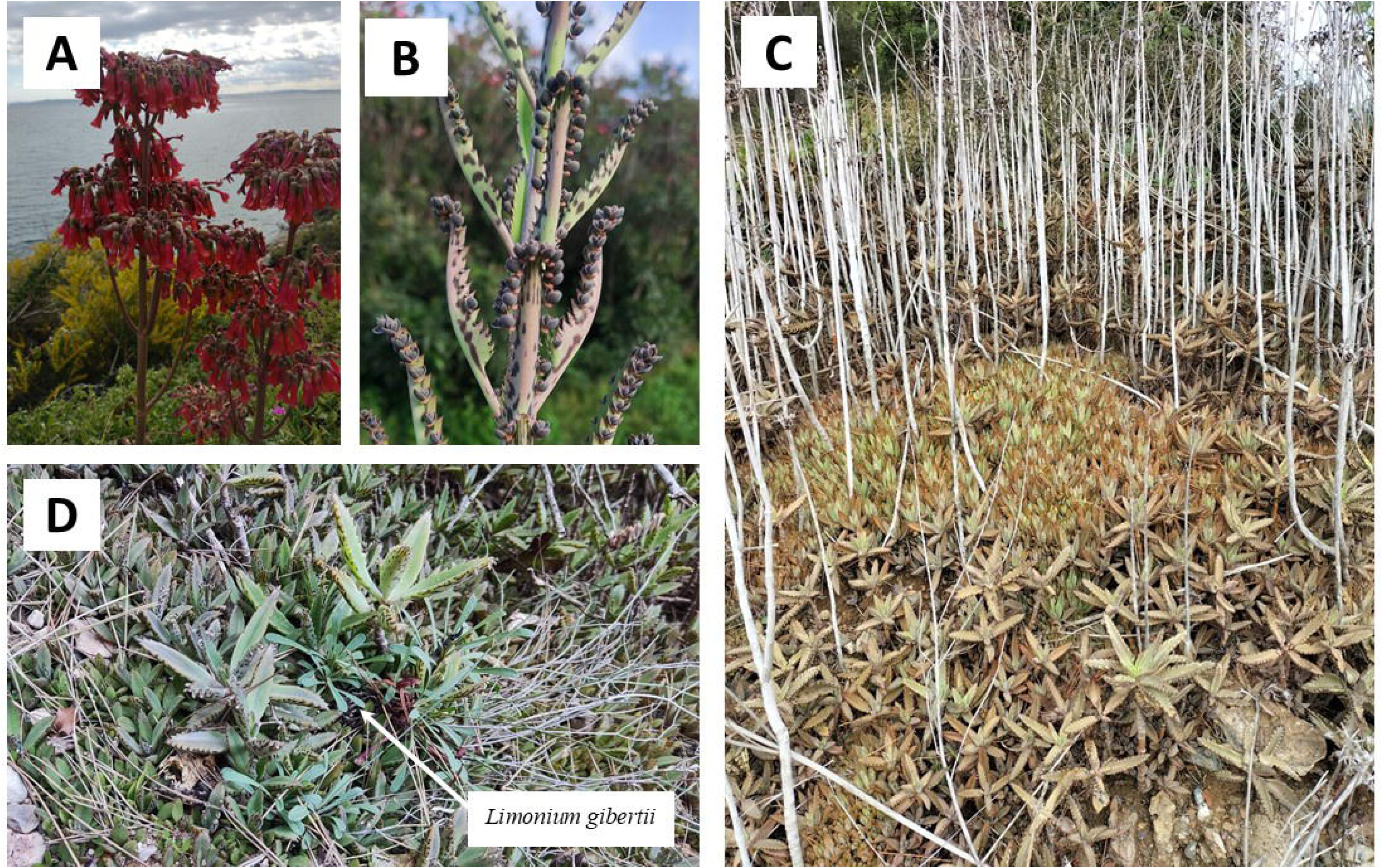
Pictures of the invader *Kalanchoe* ×*houghtonii* morphotype A. (a) pictures of the plant flowering, (b) details of adult leaves with plantlets emerging in the margins of the leaves, (c) examples of habitats invaded by *K.* ×*houghtonii*, becoming established and spreading by plantlet propagation, (d) examples of *K.* ×*houghtonii* displacing endemic *Limonium gibertii* in Montroig del Camp (Spain). Pictures credits: (a) to Jordi López-Pujol, (c) to Sònia Garcia and (b) and (d) to Joan Pere Pascual-Díaz.

Considering its likely recent origin, this complex—particularly morphotype A—exhibits an extraordinary invasive capacity. Moreover, according to preliminary species distribution models, the hybrid exhibits a larger potential distribution area compared to its parental species (Ojea-Jiménez *et al*., 2024). The invasiveness of *K.* ×*houghtonii* is granted by its successful vegetative propagation through pseudobulbils. These plantlets emerge profusely from the leaf margins (Akulova-Barlow, 2009) enabling densities up to 1,000-2,000 individuals/m^2^ when falling to the ground (Herrando-Moraira *et al*., 2020). This feature earned them the popular names of “mother of thousands” or “mother of millions” (Fig. 1). The early history of the invasion of *K.* ×*houghtonii* is poorly known, but it has reached a nearly worldwide distribution in only 80 years, being currently present in all continents except Antarctica (Herrando-Moraira *et al*., 2020). Domestic gardens are the most apparent source of incursions into the wild, at least in the US (Ward, 2006), Australia (Queensland Government 2016) and Europe (Guillot *et al*., 2014; Sáez *et al*., 2017). The first confirmed wild records of the hybrid complex were detected in the 1960s in Oceania (with the first confirmed world record in Queensland, Australia, in 1965), followed by records from North and South America in the 1970s. Then, by the 2000s, an apparent “outbreak” occurred worldwide, possibly due to a much increased detection efficiency of the hybrid thanks to the emergence of citizen science platforms (Herrando-Moraira *et al*., 2020). In the Mediterranean basin, the first naturalised record of the hybrid was reported in Calp (València, eastern Spain) in 1993 (Guillot-Ortiz *et al*., 2015; voucher VAL-930300, initially misidentified as *K. daigremontiana*).

*Kalanchoe* ×*houghtonii* poses a threat to the native plant diversity, particularly in coastal areas including some European Union habitats of conservation interest. One of the most affected habitats are the Mediterranean cliffs with endemic *Limonium* species (see Fig. 1D), given the preference for rocky substrates of *K.* ×*houghtonii* (Herrando-Moraira *et al*., 2020). *Kalanchoe* ×*houghtonii* is increasingly present in checklists of invasive flora in some Mediterranean countries, such as Algeria (Sakhraoui and Thomson, 2024) and Italy (Galasso *et al*., 2024). It also appears in the Spanish checklist of “Allochthonous species liable to compete with native wildlife, alter their genetic purity, or disrupt ecological balances” (Spanish Government, 2024). The recent appearance of this taxon in several such lists highlights the current concern of this problematic hybrid, for which management strategies are still missing. Moreover, it has been already classified as invasive in Australia (Randall, 2007; Queensland Government, 2016) and the United States (Florida Exotic Pest Plant Council, 2017).

Although several studies have explored the ecological impact of *K.* ×*houghtonii* in introduced areas (Herrera and Nassar, 2009; Herrera *et al*., 2012, 2016, 2018; Herrando-Moraira *et al*., 2020; Vargas *et al*., 2022), the genetic and genomic mechanisms underlying the invasiveness of this hybrid complex remain largely unexplored. Hybridisation can create a pool of genetic diversity, driving adaptive evolution and facilitating the emergence of novel genotypes through previously unexplored allele and gene combinations (Bock *et al*., 2016; Smith *et al*., 2020). It may also increase the structural genomic variability of taxa, including alterations in ploidy level, chromosomal rearrangements, and variations in the activity or abundance of transposable elements (McClintock, 1984). Hybridisation and polyploidy can also positively influence the invasion process—without necessarily increasing the genomic diversity—by enhancing asexual reproduction or clonality, and by increasing phenotypic plasticity of individuals (i.e. the “general-purpose genotype” strategy, Baker 1955, 1974; Bricker *et al*., 2018; Coughlan *et al*., 2017; Goad *et al*., 2021). The global invasion of *K. ×houghtonii* raises questions about the role of hybridisation and polyploidy in its high adaptability. It is unclear whether these processes contribute to increased genetic variability within the species complex or lead to the emergence of specific genotypes with higher fitness and environmental plasticity. Understanding this could shed light on how IAPS successfully adapt to diverse environments, outcompeting the native species.

Considering the extremely rapid and worldwide expansion of *K.* ×*houghtonii*, we aim to investigate the genomic variability behind its invasion success. To reach this goal, we analysed whole-genome sequencing (WGS) data together with genome size estimations, chromosome counts and ploidy levels of 57 accessions of this hybrid complex, including samples from all morphotypes and the parental species. In particular, we aim to: (1) determine the cytogenetic and genomic variability within the samples obtained from the Mediterranean basin, America and Australia through comparison between the broadly invasive morphotype A and the less invasive morphotype B; (2) disentangle the evolutionary origin of the morphotypes that occur naturally in Madagascar (i.e., morphotypes C and D); and (3) understand the relationship between specific cytogenetic and genomic features with its invasiveness.

## Materials and Methods

### Sampling and whole-genome sequencing

Samples were field-collected from representative populations of *Kalanchoe* ×*houghtonii*, *K. daigremontiana* and *K. delagoensis* across the Mediterranean basin, and stored in the living collection of the Botanical Institute of Barcelona, IBB (CSIC-CMCNB) (Table 1, Fig. 2). Sampling consisted of 30 individuals of *Kalanchoe* ×*houghtonii*, five of *K. delagoensis* and four of *K. daigremontiana*, all from different populations. Moreover, we sequenced 14 samples from herbarium vouchers across the world, including a paratype (a specimen originating from the same sampling as the holotype in Florida) and a topotype (a specimen from the type locality that also match the holotype morphologically) of *K.* ×*houghtonii*, as well as the first globally reported herbarium voucher of this hybrid (Australia). We also sampled four individuals from other *Kalanchoe* taxa (including *K.* ×*descoingsii*, *K. laetivirens*, and *K. sanctula*) closely related to *K. daigremontiana* and *K. delagoensis*. Herbarium vouchers of all specimens are deposited at the Botanical Institute of Barcelona (BC herbarium).

**FIGURE 2.**
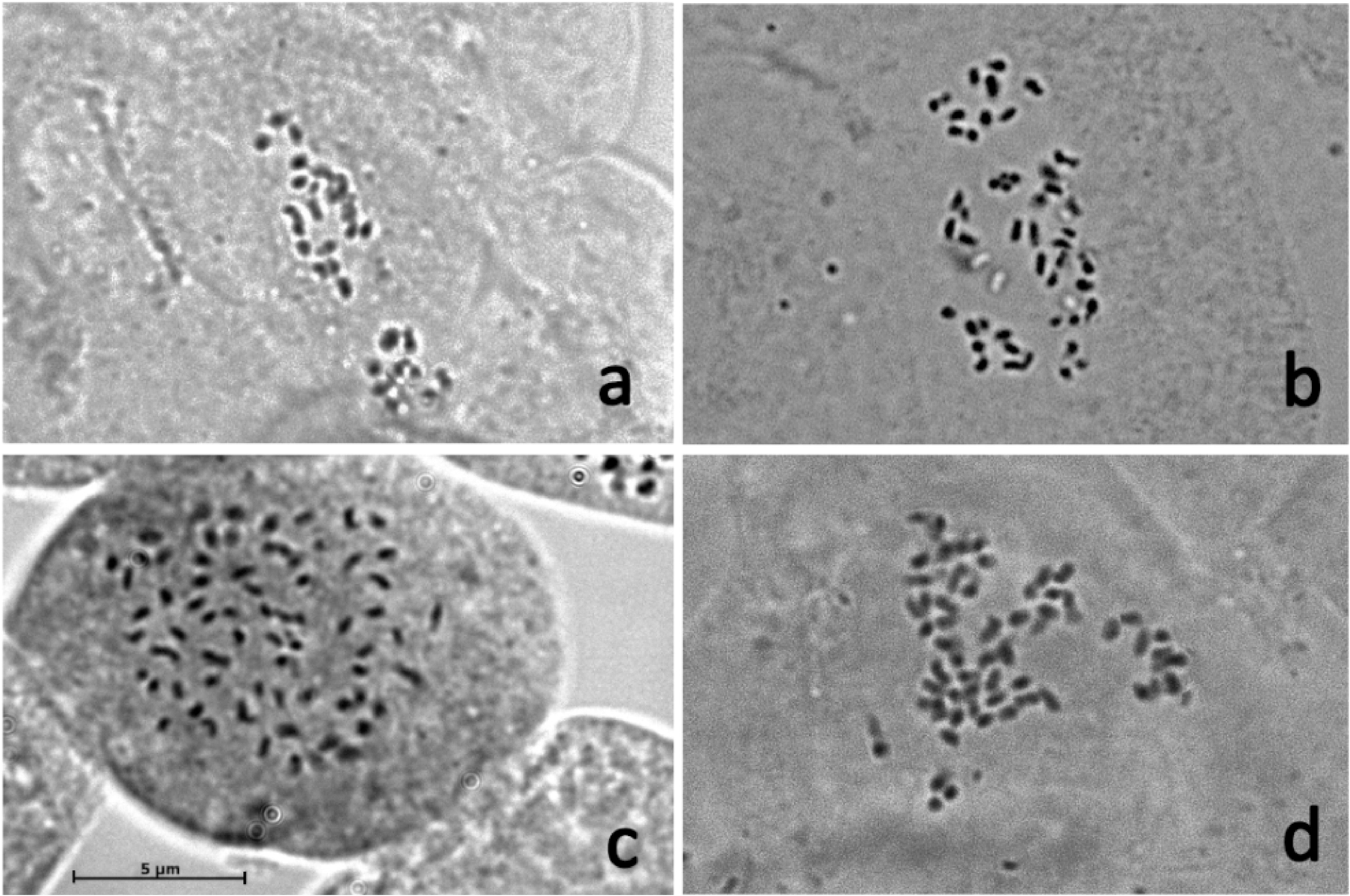
Map of the global distribution of the hybrid *Kalanchoe* ×*houghtonii* (light blue) and its parental species, *K. daigremontiana* (dark blue) and *K. delagoensis* (magenta). Data correspond to observations in GBIF from 1950 to 2023. Sampling for the present study is indicated in zoomed and coloured maps. Herbarium vouchers are indicated in orange.

**TABLE 1.** Information of all *Kalanchoe* populations sampled. Paratype (*) and topotype (**) vouchers are indicated. KH indicates *Kalanchoe* ×*houghtonii*, KDA indicates *Kalanchoe daigremontiana*, KDE indicates *Kalanchoe delagoensis*.

| Sample | Morphotype | Wild (W) /<br>Cultivar (C) | Location | Year |
| --- | --- | --- | --- | --- |
| KDA (Bahamas) | Parental | W | Bimini, Bahamas | 1975 |
| KDA (Israel 3) | Parental | W | Haifa, Israel | 2022 |
| KDA (Israel 2) | Parental | W | Tel Aviv, Israel | 2020 |
| KDA (Tunisia 2) | Parental | W | Tunis Bizerte, Tunisia | 2019 |
| KDA (Spain 1) | Parental | W | València, Spain | 2019 |
| KDE (Tunisia 1) | Parental | W | Le Kram, Tunisia | 2019 |
| KDE (Madagascar 2) | Parental | W | Behara, Madagascar | 2022 |
| KDE (Madagascar 1) | Parental | W | Antananarivo, Madagascar | 2022 |
| KDE (Israel 1) | Parental | W | Tel Aviv, Israel | 2020 |
| KDE (Catalonia 1) | Parental | W | Tossa de Mar, Catalonia | 2019 |
| KH (Australia 1) | A | W | Chinchilla, Australia | 1970 |
| KH (Australia 2) | A | W | Ipswich, Australia | 1970 |
| KH (Australia 3) | A | W | Ipswich, Australia | 1966 |
| KH (Florida 1) | A | W | Monroe Co., Florida, US | 2009 |
| KH (Florida 2)* | A | W | Fort Myers Beach, Florida, US | 2000 |
| KH (Florida 3)** | A | W | Merritt Island, Florida, US | 2004 |
| KH (Ecuador) | A | W | Mitad del Mundo, Ecuador | 1988 |
| KH (Venezuela 2) | A | W | Churuguara, Venezuela | 1980 |
| KH (Dominican Republic) | A | W | San José de Ocoa, Dominican Republic | 1993 |
| KH (Spain 5) | A | W | Calp, Spain | 1993 |
| KH (Catalonia 4) | A | W | Garraf, Catalonia, Spain | 2003 |
| KH (Greece) | A | W | Athens, Greece | 2019 |
| KH (France 2) | A | W | Banyuls, France | 2023 |
| KH (Italy 3) | A | W | Camerota, Italy | 2023 |
| KH (Morocco) | A | W | Casablanca, Morocco | 2019 |
| KH (“Palamós”) | A | C | In cultivation | - |
| KH (Italy 2) | A | W | Formia, Italy | 2023 |
| KH (Spain 2) | A | W | Almeria, Spain | 2019 |
| KH (Italy 4) | A | W | Genova, Italy | 2023 |
| KH (‘Jaws of Life’) | A | W | Cultivar | - |
| KH (Italy 7) | A | W | Lipari, Aeolian Islands, Italy | 2023 |
| KH (Portugal) | A | W | Lisboa, Portugal | 2018 |
| KH (Italy 8) | A | W | Livorno, Italy | 2023 |
| KH (Malta) | A | W | Mosta, Malta | 2019 |
| KH (France 1) | A | W | Cap d’Ail, France | 2023 |
| KH (Italy 5) | A | W | Ostia, Italy | 2023 |
| KH (Spain 3) | A | W | Alacant, Spain | 2019 |
| KH (Italy 6) | A | W | La Maddalena, Sardegna, Italy | 2023 |
| KH (Catalonia 3) | A | W | Tarragona, Catalonia, Spain | 2023 |
| KH (France 3) | A | W | Toulon, France | 2023 |
| KH (Spain 4) | A | W | Tórrox, Spain | 2022 |
| KH (Tunisia 2) | A | W | Tunis Bizerte, Tunisia | 2019 |
| KH (Australia 4) | B | W | Byrnestown, Australia | 1965 |
| KH (Australia 5) | B | W | Gracemere, Australia | 1968 |
| KH ('Pink Butterflies') | B | C | Cultivar | - |
| KH (Catalonia 2) | B | W | Cap de Creus, Catalonia, Spain | 2021 |
| KH ('Hybrida') | B | C | Cultivar | - |
| KH (Italy 1) | B | W | Ostia, Italy | 2023 |
| KH ('Linear leafed') | C | C | Cultivar | - |
| KH (Madagascar 3) | D | W | Makay Massif, Madagascar | 2022 |
| KH (Madagascar 4) | D | W | Tulear, Madagascar | 2022 |
| KH ('Parsel Tongue') | D | C | Cultivar | - |
| *Kalanchoe* ("RS574") | - | C | Cultivar | - |
| *Kalanchoe laetivirens* | - | W | Tulear, Madagascar | 2022 |
| *Kalanchoe sanctula* | - | W | Taolagnaro, Madagascar | 2022 |
| *Kalanchoe* ×*descoingsii* | - | W | Arboretum Antsokay, Madagascar | 2022 |

Total DNA was isolated from fresh leaves using two different modified CTAB protocols, one general protocol for fresh leaves material (Doyle and Doyle, 1987) and another for samples from ancient herbarium vouchers (Hale *et al*., 2020). The quality of each sample was checked by spectrophotometry with NanoDrop 1000 (PeqLab, Erlangen, Germany) and the DNA concentration by fluorometry with Qubit Fluorometric Quantification (Thermo Fisher Scientific, Waltham, MA, US). The genomic DNA was randomly sheared into short fragments and sequenced by NovoGene Europe (Cambridge, UK). Libraries of the whole genome with an average insert size of 450 bp were sequenced on an Illumina NovaSeq Platform (Illumina, San Diego, CA, US). For all accessions, around 2.68–13.3 Gbp of raw data (equivalent to ∼10× of their respective genome size) were generated with pair-end 150-nt read length. We generated 375 Gb of whole-genome sequencing data for 57 *Kalanchoe* accessions (with one sample being replicated to determine the batch sequencing error in downstream analyses), with an emphasis on sampling *Kalanchoe ×houghtonii* from across its range in the Mediterranean basin (*n* = 20; Fig. 2).

### Ploidy level and genome size estimation using flow cytometry

Ploidy level and genome size estimations were carried out with a flow cytometer CyFlow® Space (Sysmex-Partec, Norderstedt, Germany), coupled with the FloMax software (Partec GmbH, Münster, Germany). The internal standards used were *Petunia hybrida* cv. ‘PxPc6’ (obtained from the living collection at the greenhouses of the IBB), with a 2C genome size of 2.85 pg, and *Solanum lycopersicum* cv. ‘Stupické polní rané’, with a 2C genome size of 1.96 pg (Temsch *et al*., 2022). Fresh young leaves were chopped using a razor blade in the GPB buffer (Loureiro *et al*., 2007) and stained by adding 40 μL of 1 mg mL^−1^ propidium iodide solution. Nuclei suspensions were then incubated for ca. 20 min on ice prior to analysis. We measured three replicates for each population, with a minimum of 500 nuclei per fluorescence peak in each analysis.

### Karyological observations

For the karyological observations, root tips of *K. ×houghtonii*, *K. daigremontiana* and *K. delagoensis* were sampled from adult plants early in the morning, prior to generous watering of the adult plant one or two days earlier. Root tips were pre-treated with ice-cold water for 24 h, following recommendations for species with very small chromosomes and then fixed in Farmer’s solution (3:1 ratio of absolute pure ethanol and glacial acetic acid). After fixation, acid hydrolysis with 1N HCl at 60°C for 10–15 minutes, followed by staining with 1% acetic orcein (for at least one hour), was conducted. Subsequently, individual root tips were selected with a magnifying glass, and the radical meristem was cut, discarding the rest of the tissue. The sample was then placed on a slide with a drop of 45% acetic acid/glycerine (9:1) solution, squashed with tweezers or a scalpel and covered with a coverslip, following Olanj *et al*. (2015). Finally, preparations were observed under the optical microscope (Zeiss Axioplan) and photographed with a coupled AxioCam HRm camera.

### Plastome assembly and phylogenomics

Plastomes were assembled from raw sequencing data, using NOVOPlasty v. 4.2.1 (Dierckxsens *et al*., 2017) with default parameters and previously removing adapters Illumina reads, as recommended by the author. Chloroplast genome assemblies were done using as seed the *rbc*L gene of *K. daigremontiana* (NCBI GenBank: L11189) and, if the assembly was incomplete, the process was repeated using a different seed sequence (the *mat*K gene, NCBI GenBank: AF274619, or the entire chloroplast sequence of *K. daigremontiana*, NCBI GenBank: MT954417). The resulting contig options were aligned to the pre-existing *K. daigremontiana* plastome genome (NCBI GenBank: MT954417) using MAFFT (Katoh and Standley, 2013) and manually rearranged so that the short single copy and inverted repeat were in the same orientation for each individual using Geneious Prime v. 2023.2.1 (Kearse *et al*., 2012). Then, filtered whole genome sequencing reads with a minimum quality score of 30 were mapped to the consensus plastome sequence using *bwa* (Li and Durbin, 2009), and the resulting consensus, with a minimum coverage of 20 reads per site, was manually adjusted for further phylogenetic analyses. All sites not supported by 90% of the mapped reads were replaced by Ns. No ambiguities were allowed since the inheritance of the plastome is determined by only one parental individual. For four herbarium accessions where the quality of reads was not good enough to allow the *de novo* reconstruction of the plastome, a reference-based reconstruction was performed using reads with a minimum quality score of 30 (using *fastp* pipeline, Chen *et al*., 2018). Herbarium voucher Australia 1 was excluded from the phylogenetic analysis due to their high proportion of missing data (Table S2). Each resulting plastome was annotated using the software GeSeq (Tillich *et al*., 2017) included in the platform MPI-MP CHLOROBOX (https://chlorobox.mpimp-golm.mpg.de/, accessed 11 February 2024), selecting the options to perform ARAGORN v. 1.2.38 and BLAT, as recommended by authors for plastomes.

Plastome phylogenomics were inferred without one of the inverted repeats, because the inverted repeats within a plastome can recombine with each other, they are generally identical, and including them effectively inflates the weight assigned to those positions (Blowers *et al*., 1989, Palmer, 1985). Plastomes were subsequently partitioned based on their tRNA, rRNA, CDS (separate introns and exons), and non-coding regions. The resulting sequences were aligned using MAFFT (Katoh and Standley, 2013). Then, we used PartitionFinder2 (Lanfear *et al*., 2017) to fit the best nucleotide substitution model for all the different partitions. Bayesian inference (BI) phylogenomic tree was conducted using the software MrBayes v. 3.2.6 (Ronquist *et al*., 2012), where two independent Markov chains Monte Carlo (MCMC) were run for 3,000,000 generations, with tree sampling every 1000 generations. The average standard deviation was confirmed to be less than 0.01, and the potential scale reduction factor was near 1.0 in all parameters. The first 25% of the trees were discarded as “burn-in”, and the posterior probability was estimated by constructing the 50% majority-rule consensus tree. Phylogenomic trees were visualised with FigTree v. 1.4.5 (Rambaut *et al*., 2024), using as outgroup *Kalanchoe humifica*, according to Rodewald *et al*. (2025).

### Ribosomal DNA reconstruction through TAREAN pipeline

Ribosomal DNA (rDNA) identification and reconstruction by similarity-based clustering of Illumina paired-end reads was performed following the TAREAN pipeline (Novak *et al*., 2017). First, Illumina FASTQ files were filtered to avoid the presence of adapters, reads with indeterminate bases (N) and a minimum quality score of 30 using the *fastp* pipeline. After converting filtered FASTQ reads to interlaced FASTA files, clustering analyses were performed on these data using the following settings: minimum overlap = 55 and cluster size threshold = 0.01%. Chloroplast and mitochondrial reads were removed before downstream analyses. For chloroplast reads, we mapped the filtered reads to the previously reconstructed plastome sequences. For mitochondrial reads, we mapped filtered reads of all accessions to *Sedum plumbizincicola* mitochondrial genome (NCBI GenBank: OP588116). The total number of reads used as input for the individual clustering analyses corresponds to the 0.5× of the genome coverage for each accession. Then, individual TAREAN analyses were carried out for each accession, allowing the cluster merging (--merge_threshold 0.1) and the automatic filtering of abundant satellite repeats (--automatic_filtering) to allow more reads to be analysed and increase the chances of reconstructing the rDNA. For the 35S and 5S rDNA phylogenetic network analyses, we extracted the consensus sequence identified in the TAREAN analyses. Once all consensus sequences were obtained, the different regions constituting the rDNA units were determined by BLAST using as reference the 35S (NCBI GenBank: X52322) and 5S (NCBI GenBank: ATHRR5S) rDNA genes of *Arabidopsis thaliana*. Afterward, we mapped the quality trimmed Illumina reads from each sample to their respective 35S and 5S rDNA consensus sequences using *bwa*. We considered the potential intra-genomic variability allowing the presence of ambiguous bases (>10% of the reads) in the final consensus sequence. For the 35S rDNA arrays, genes were discarded for downstream analysis since there was presence of DNA contamination from other organisms (e.g. fungi, human DNA) likely due to the poor DNA preservation in some herbarium vouchers. Consensus sequences of all samples were first aligned using MAFFT, and then we reconstructed a Neighbor-net (Bryant and Moulton, 2004) by transforming sequence divergence to uncorrected P-distances and handling ambiguous characters as average states using SPLITSTREE (Huson and Bryant, 2024).

### Nuclear phylogenomics based on BUSCO genes

To better understand the hybridisation process and the genetic diversity exhibited by *K. ×houghtonii*, we examined the composition of nuclear polymorphisms using Benchmarking Universal Single-Copy Ortholog genes (BUSCO, Seppey *et al*., 2019). The list of BUSCO genes used was obtained from the *Kalanchoe fedtschenkoi* partially assembled genome (*Kalanchoe fedtschenkoi* v. 1.1, Yang *et al*., 2017), setting eudicots_odb10 as the lineage database. The resulting list of 1,896 BUSCO genes were used as targets for bait sequence using HybPiper (Johnson *et al*., 2016). For each sample, we filtered each one of the BUSCO genes that hold three or more paralogs using the implemented HybPiper script “paralogs_retriever.py”, to avoid excessive heterozygosity and coverage, for both polyploid and diploid samples (Bohutínská *et al*., 2023). According to HybPiper statistics (reads mapped and gene recovery), populations Australia 1, Australia 2, Australia 3, Ecuador, Venezuela 2 and Dominican Republic were discarded from downstream analyses (Table S3). Then, following Pokorny *et al*., (2024), the max_overlap.R script (Shee *et al*., 2020) was used to identify underrepresented, incomplete, and unevenly distributed sequences, where genes with < ⅔ median coverage score values (computed by the above mentioned script) were also discarded from downstream analyses.

MAFFT was used to generate alignments for individual BUSCO genes using the default parameters. Multiple sequence alignment (MSA) summary statistics were computed then with AMAS (Borowiec 2016) to assess quality. BUSCO genes were excluded if the number of taxa, or the proportion of parsimony informative sites (P_PIS_) was < ⅓ of the median value across all genes, or if the percentage of missing data was > ⅓ of the median value across all genes. Resulting MSAs were used to infer exploratory trees with FastTree2 (Price *et al*., 2010) for automated outlier removal with TreeShrink (Mai and Mirarab 2018) in “per-species” mode for various levels of false positive tolerance (α), which controls outlier detection (-q “0.01 0.05 0.5”). Pre- and post-automated outlier removal FastTrees were visually inspected with Geneious Prime to check TreeShrink performance. Outlier-filtered data matrices (0.01 threshold) were realigned (using MAFFT), and summary statistics were computed as above (keeping a total number of 1,122 filtered BUSCO genes). Output MSAs were refined with trimAl (Capella-Gutiérrez *et al*., 2009), using lax gap and conservation thresholds (-gt 0.3 - cons 30) to prevent the massive loss of data and phylogenetic signal (P_PIS_).

Gene trees for each one of the remaining BUSCO genes were estimated with IQ-TREE 1.5.5 (Nguyen *et al*., 2015) using ModelFinder Plus (Kalyaanamoorthy *et al*., 2017), to select the best-fit model and continued with maximum likelihood (ML) tree inference and using both UFBoot (Ultrafast bootstrap, Minh *et al*., 2013) and SH-like (Guindon *et al*., 2010) approximation, to compute 1,000 bootstrap replicates. The resulting gene trees had bipartitions collapsed under various bootstrap support (BS) thresholds (’i & b<’$bs’’) using the nw_ed pipeline from the newick_utils set of programs (Junier and Zdobnov, 2010). These variously collapsed BUSCO gene trees were used as input to estimate the nuclear-based species trees with ASTRAL III 5.6.3 (Zhang *et al*., 2018), which was run with extensive Newick annotations (-t 2). As recommended by Zhang *et al*., (2018), the selected final tree was the one showing collapsing bipartitions with extremely low support (‘i & b<0’) since this strategy can substantially improve accuracy.

### SNP filtering and variant analyses

Good quality Illumina reads were mapped to each of the filtered BUSCO genes for every population with good gene recovery using *bwa*. Following Bohutínská *et al*., (2023), Genome Analysis Toolkit (GATK, McKenna *et al*., 2010) was used to remove duplicated sequences or PCR technical errors (MarkDuplicates) which can inflate the sequencing depth. Then, two separate variant-calling (HaplotypeCaller) datasets were conducted for each population: one considering the ploidy level (using –ploidy option), and one without considering it. Then, within each dataset, variant call files were combined (CombineGVCFs) and genotyped (GenotypeGVCFs), reaching a total number of 148,908 unfiltered single nucleotide polymorphisms (SNPs) for the non-ploidy dataset and 34,893 SNPs for the ploidy dataset. BCFTOOLS version 1.11 (Li, 2011) was used to remove SNPs within 20 bp of an indel or other variant type, keeping only bi-allelic SNPs (72,684 SNPs for non-ploidy dataset and 24,009 SNPs for the ploidy dataset). Then, SNPs were marked by VariantFiltration and excluded with SelectVariants if they meet with one or more of the following criteria: (i) Quality by Depth (QD) < 10.0, (ii) Fisher Strand Bias (FS) > 60.0, (iii) Strand Odds Ratio (SOR) > 3.0, (iv) RMS Mapping Quality (MQ) < 40, (v) -2.5 < Mapping Quality Rank Sum Test (MQRankSum) > 2.5, (vi) Depth (DP) > 1426.2 (two times average DP) and (vii) Read Position Rank Sum Test (ReadPosRankSum) > 2.5. Then VCFTOOLS version 0.1.15 (Danecek *et al*., 2011) was used to convert individual genotypes to missing data when the genotype quality (--minGQ) was lower than 30 and the depth of coverage (--minDP) was lower than 10. Ultimately, the filtering resulted in 44,055 SNPs for the non-ploidy dataset and 9,763 SNPs for the ploidy dataset.

The non-ploidy dataset of SNPs was used to perform a principal component analysis (PCA). For this, PLINK (Purcell *et al*., 2007) was used to generate eigenvectors and eigenvalues, previously pruning SNPs to remove high linkage disequilibrium (LD) (-- indep-pairwise 50 5 0.7), keeping a total of 11,966 SNPs. Then, PC1, PC2 and PC3 (Fig. 5C, Fig. S2) outputs were visualised using the R package *ggplot2* (Wickham, 2016). Moreover, the non-ploidy dataset was used to assess the clonality of populations from morphotypes A, B, and D (excluding C since only one population for this morphotype is available). First, the shared heterozygosity (SH) method (Yu *et al*., 2022) was used to identify clonemates, based on the number of heterozygous sites relative to the number of heterozygous sites observed in the individual with the higher count (Fig. 5A). Furthermore, Identity-By-Descent (IBD) was also used to define the clonal relationship for pairwise comparisons among individuals (Fig. 5B). The IBD analysis calculated the proportion of the SNPs with zero, one or two shared IBD alleles to evaluate genetic similarity among individuals. For this, PLINK was used to calculate IBD values for pair-wise comparisons among individuals ––previously pruning SNPs to remove high LD (--indep-pairwise 50 5 0.7)–– and we considered pairs of individuals to be a clonal relationship if they had an IBD value greater than 0.95 (Bonfante *et al*., 2021; Cong *et al*., 2022; Liang *et al*., 2019; Migicovsky *et al*., 2017; Myles *et al*., 2011; Zou *et al*., 2023).

The ploidy dataset of SNPs was used to identify nuclear genetic clusters of the different morphotypes using ENTROPY version 2.0, designed for quantifying population structure in autopolyploid and mixed-ploidy individuals using genotype-likelihood data (Shastry *et al*., 2021). As recommended by authors, for each genetic cluster (*K*) we spawned three simultaneous MCMC to assess convergence. The maximum *K* in the genotype likelihoods was 6 (i.e. the sum of parental species and hybrid morphotypes), and the optimal *K* was determined by the deviance information criterion (DIC) and the log-pointwise predictive density (Table S6).

Finally, the intragenomic polymorphisms present in the nuclear data were analysed to assess the hybridisation and gene flow between samples. For this, we generated consensus sequences for all filtered BUSCO genes by mapping (with *bwa* v. 0.0.17) the good quality FASTQ files to the selected BUSCO genes with good coverage and with low missing data (total of 1,122 genes). Then, consensus sequences were aligned and refined using MAFFT and trimAl, respectively, obtaining a final alignment of 4,000,577 bp. The obtained matrix was used as input for SPLITSTREE, reconstructing a Neighbor-net, transforming sequence divergence to uncorrected P-distances, and handling ambiguous characters as average state.

## Results

### Ploidy level and genome size estimations from flow cytometry measurements and chromosome counts

The measurements of nuclear genome size (2C values) and ploidy level estimation were conducted on 42 samples (Table 2), comprising four *K. daigremontiana*, five *K. delagoensis*, and 33 *K. ×houghtonii* populations. In all samples, endopolyploidy was common, depending on the plant material used for the measurements. *Kalanchoe daigremontiana* exhibited a genome size ranging from 0.54 to 0.58 pg, corresponding to a diploid ploidy level. *Kalanchoe delagoensis* had a genome size ranging from 1.05 to 1.16 pg, corresponding to a tetraploid ploidy level. For *K. ×houghtonii* hybrid complex, we found three different ploidy levels: (i) the morphotype A had a genome size ranging from 1.04 to 1.18 pg, corresponding to a tetraploid, (ii) the morphotypes B and C had a genome size ranging from 0.79 to 0.85 pg, corresponding to a triploid, and (iii) morphotype D had a genome size ranging from 0.53 to 0.59 pg corresponding to a diploid. Chromosome counts confirmed 2*n* = 2*x* = 34 for *K. daigremontiana* (population “Israel 2”), 2*n* = 4*x* = 68 for *K. delagoensis* (population “Catalonia 1”), 2*n* = 3*x* = 51 for *K. ×houghtonii* morphotype B (population “Catalonia 2”), and 2*n* = 4*x* = 68 for *K. ×houghtonii* morphotype A (population “Palamós”). Pictures are shown in Fig. S3.

**TABLE 2.** List of all *Kalanchoe* populations whose genome size and ploidy level estimations have been analysed. The amount of DNA is expressed in 2C-value and picograms. KH indicates *Kalanchoe* ×*houghtonii*, KDA indicates *Kalanchoe daigremontiana*, KDE indicates *Kalanchoe delagoensis*.

| Sample | Genome size (pg) | CV (%) | Ploidy level estimation |
| --- | --- | --- | --- |
| KDA (Israel 3) | 0.56 | 13.53 | Diploid |
| KDA (Israel 2) | 0.58 | 11.31 | Diploid |
| KDA (Tunisia 2) | 0.54 | 11.57 | Diploid |
| KDA (Spain 1) | 0.58 | 12.39 | Diploid |
| KDE (Tunisia 1) | 1.16 | 9.37 | Tetraploid |
| KDE (Madagascar 2) | 1.08 | 10.16 | Tetraploid |
| KDE (Madagascar 1) | 1.05 | 9.15 | Tetraploid |
| KDE (Israel 1) | 1.06 | 12.99 | Tetraploid |
| KDE (Catalonia 1) | 1.10 | 8.92 | Tetraploid |
| KH (Greece) | 1.05 | 7.71 | Tetraploid |
| KH (France 2) | 1.08 | 10.80 | Tetraploid |
| KH (Italy 3) | 1.04 | 18.00 | Tetraploid |
| KH (Morocco) | 1.08 | 10.96 | Tetraploid |
| KH (“Palamós”) | 1.14 | 9.43 | Tetraploid |
| KH (Italy 2) | 1.06 | 8.81 | Tetraploid |
| KH (Spain 2) | 1.07 | 6.51 | Tetraploid |
| KH (Italy 4) | 1.14 | 10.67 | Tetraploid |
| KH (‘Jaws of Life’) | 1.12 | 5.89 | Tetraploid |
| KH (Italy 7) | 1.04 | 6.94 | Tetraploid |
| KH (Portugal) | 1.10 | 6.58 | Tetraploid |
| KH (Italy 8) | 1.13 | 4.69 | Tetraploid |
| KH (Malta) | 1.13 | 5.63 | Tetraploid |
| KH (France 1) | 1.13 | 9.20 | Tetraploid |
| KH (Italy 5) | 1.13 | 5.63 | Tetraploid |
| KH (Spain 3) | 1.17 | 8.43 | Tetraploid |
| KH (Italy 6) | 1.17 | 8.03 | Tetraploid |
| KH (Catalonia 3) | 1.15 | 8.49 | Tetraploid |
| KH (France 3) | 1.12 | 4.88 | Tetraploid |
| KH (Spain 4) | 1.18 | 8.32 | Tetraploid |
| KH (Tunisia 2) | 1.10 | 8.24 | Tetraploid |
| KH (‘Pink Butterflies’) | 0.85 | 8.15 | Triploid |
| KH (Catalonia 2) | 0.79 | 11.28 | Triploid |
| KH (Italy 1) | 0.83 | 11.94 | Triploid |
| KH (‘Hybrida’) | 0.84 | 7.42 | Triploid |
| KH (‘Linear leafed’) | 0.84 | 5.41 | Triploid |
| KH (Madagascar 3) | 0.59 | 12.31 | Diploid |
| KH (Madagascar 4) | 0.58 | 6.30 | Diploid |
| KH (‘Parsel Tongue’) | 0.53 | 9.60 | Diploid |
| *Kalanchoe* (“RS574”) | 0.80 | 5.61 | Triploid |
| *Kalanchoe laetivirens* | 1.59 | 3.91 | Hexaploid |
| *Kalanchoe sanctula* | 0.57 | 6.67 | Diploid |
| *Kalanchoe xdescoingsii* | 1.30 | 3.74 | Pentaploid |

### Phylogenetic relationships inferred from whole plastome sequence analysis

The whole plastome was completely reconstructed for all accessions, including those from old herbarium vouchers. The length of the plastome sequences differed between accessions: all *K. delagoensis* accessions and the morphotype C (3*x*) of *K. ×houghtonii* were 150,018 bp long except for the *K. delagoensis* population “Madagascar 1”, which had a length of 149,975 bp; all *K. daigremontiana* accessions and the morphotype A (4*x*) of *K. ×houghtonii* were 150,062 bp long; and all morphotype B accessions of *K. ×houghtonii* (3*x*) were 150,056 bp long. Morphotype D and the other *Kalanchoe* taxa studied exhibited a range of lengths between 149,923 and 150,173 bp (see Table S2 for further details).

The analysis of phylogenetic relationships based on plastome sequences (124,637 bp alignment, with 85.8% identical sites after removal of one of the inverted repeats) is depicted in Fig. 3. The whole plastome tree is rooted to *K. humifica*, and an early diverging major clade is constituted by all samples of *K. delagoensis*, as well as those of *K. ×houghtonii* morphotype C and D. In this clade, *K. ×houghtonii* “Madagascar 3” is the first splitting sample, followed by the subclade constituted by *K. ×houghtonii* ‘Parsel Tongue’ and “Madagascar 4” (i.e. samples of morphotype D). All *K. delagoensis* and *K. ×houghtonii* ‘Linear leafed’ (i.e. morphotype C) samples constitute another subclade, where *K. delagoensis* “Madagascar 1” diverges first, and the rest of samples together form a polytomy (*K. delagoensis* “Catalonia 1”, “Israel 1”, “Madagascar 2”, “Tunisia 1” and *K. ×houghtonii* ‘Linear leafed’). The other major clade of the tree is constituted by samples of *K. sanctula*, *Kalanchoe* “RS574”, *K. daigremontiana*, *K. laetivirens*, *K*. ×*descoingsii* and *K*. ×*houghtonii* morphotypes A and B. This major clade splits in two further clades: one constituted by *K. sanctula* and *Kalanchoe* “RS574”, appearing as the sister clade of a second larger clade consisting of all samples of *K. daigremontiana, K. laetivirens, K. ×descoingsii* and those of *K. ×houghtonii* morphotypes A and B. Three main subclades are nested within this second clade, one including *K. laetivirens* and *K. ×descoingsii* samples, the second including all *K. ×houghtonii* morphotype B samples, and the third encompassing all sampled populations of *K. daigremontiana* and all of *K. ×houghtonii* morphotype A. The morphotype A and *K. daigremontiana* subclade includes all samples collected in the Mediterranean basin identified as morphotype A and historical vouchers such as the paratype and the topotype of *K. ×houghtonii* from Florida (USA), the first record of this hybrid in the Mediterranean region, and other herbarium accessions from Australia, Ecuador, Bahamas, Florida, Venezuela, and the Dominican Republic, all together forming a polytomy in exception of the population Australia 3, which is slightly separated from the rest without support (pp = 0.93). The morphotype B subclade, which includes populations from Spain (population from Cap de Creus, Catalonia), Italy, herbarium accessions from Australia (including the first globally reported herbarium voucher identified as *K.* ×*houghtonii*) and the cultivars ‘Hybrida’ and ‘Pink butterflies’, also forms a polytomy.

**FIGURE 3.**
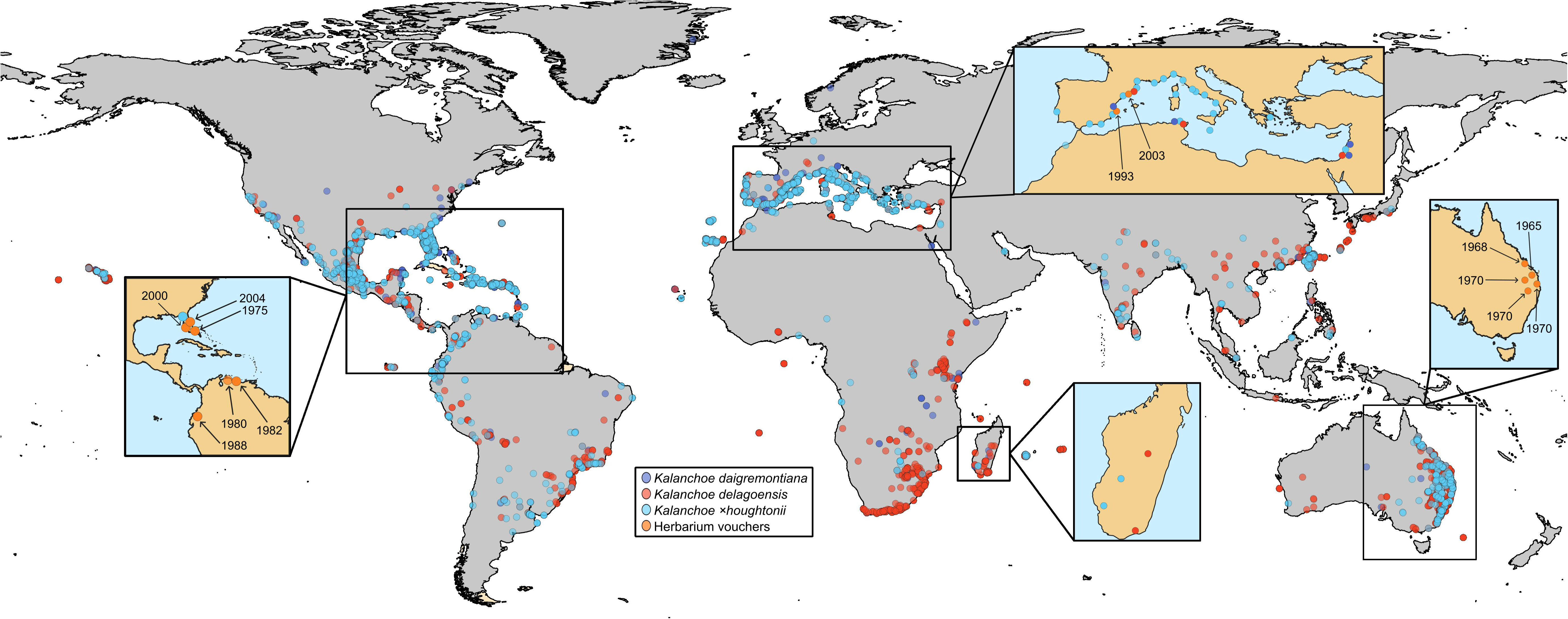
Phylogenomic relationships of 58 samples of the *Kalanchoe* ×*houghtonii* hybrid complex based on 125,386 bp of the whole plastid genome (with one inverted repeat removed) using Bayesian Inference approach. Samples of *Kalanchoe* ×*houghtonii* morphotype A are indicated in light blue, morphotype B in purple, morphotype C in yellow and morphotype D in orange. Samples of parental species are indicated in dark blue for *Kalanchoe daigremontiana* and magenta for *Kalanchoe delagoensis*. Nodes without support values are considered maximum supported (pp = 1). Technical replicates are indicated with superindex 1 and 2. The whole plastome sequence of *Kalanchoe humifica* is used as an outgroup. Pictures credits: KDA to Robin White, KDE to Steve K., KH morphotype A to Piambr, morphotype B to Chris Bentley, morphotype C and morphotype D to Solofo E. Rakotoarisoa.

### Phylogenetic relationships inferred from ribosomal DNA

The phylogenetic relationships among samples of *K. ×houghtonii* complex based on 35S and 5S rDNA sequences are presented in Fig. S1. The alignment of 35S rDNA sequences consisted of 3574 bp, excluding the 18S, 5.8S, and 26S genes. For the 5S rDNA, 467 bp were aligned, including the 5S gene and the non-transcribed spacer (NTS). For the 35S rDNA, 96.0% of sites were identical, while for the 5S rDNA, 86.3% of sites were identical.

Both 35S and 5S rDNA networks exhibited the same overall topology. Parental accessions (*K. daigremontiana* and *K. delagoensis*) were distanced between them, without sharing direct reticulations. For both 35S and 5S rDNA networks, all samples from *K. daigremontiana* were grouped, while for *K. delagoensis* all samples were grouped except for one population from Madagascar, which was more closely related to the hybrid morphotype A in the 35S rDNA network. Samples from morphotypes A and B were closely related between them and, in the 5S rDNA network, morphotype B nested inside morphotype A cluster. Morphotype C also shared reticulations with both parental species, but the extent of connections differed between 5S and 35S rDNA networks. For the 35S rDNA, morphotype C was more connected to *K. daigremontiana*, whereas for the 5S rDNA it was closer to *K. delagoensis*. Finally, morphotype D was more closely related to the *K. daigremontiana* cluster, being its closest morphotype in both 35S and 5S rDNA networks.

Regarding the other *Kalanchoe* taxa (K. ×*descoingsii*, *K. laetivirens*, and *K. sanctula*), both 5S and 35S networks exhibited similar topologies. In the 35S rDNA network, all these other taxa were placed into a new branch closely related to *K. delagoensis* samples, with long branches. In contrast, for the 5S rDNA both *K. ×descoingsii* and *K. laetivirens* were more related with morphotype A and *K. daigremontiana*, but *K. sanctula* was separated by a long branch from the rest of studied taxa. In contrast, the unidentified *Kalanchoe* sample ‘RS574’ was closely related to the *K. delagoensis* group in both 35S and 5S rDNA.

### Nuclear genetic variation and structure within *Kalanchoe ×houghtonii*

The phylogenomic relationships based on nuclear DNA were inferred using data from 1,122 BUSCO genes across 47 populations (Fig. S4). The tree is rooted to *K. humifica*, with *K. sanctula* positioned as the earliest-divergent species. The first major clade contains *Kalanchoe* “RS574” diverging in a first branch from a subclade constituted by *K. delagoensis* and *K. ×houghtonii* ‘Linear leafed’ (i.e. morphotype C). Within this subclade, *K. delagoensis* population “Madagascar 1” diverges first, followed by *K. ×houghtonii* ‘Linear leafed’ and the remaining *K. delagoensis* populations that are grouped together. In the other major clade of the tree, *K. ×descoingsii* appears in a sister position to the rest of the samples, clustered in two large subclades. The first of these subclades encompasses all the samples from *K. ×houghtonii* morphotype B, while the other large subclade includes samples from *K laetivirens*, *K. ×houghtonii* morphotype D, *K. daigremontiana* and *K. ×houghtonii* morphotype A. Finally, this last subclade splits into two distinct groups: (i) a group containing all populations of *K. daigremontiana* and *K. ×houghtonii* morphotype D, with *K. laetivirens* placed in a sister position to morphotype D samples, and (ii) a group containing all *K*. *×houghtonii* morphotype A populations, except for “Florida 1”, which is separated from the rest of the morphotype A without support (support = 0.52).

According to the analysis of population structure (ENTROPY), the optimal number of genetic clusters (*K*) for all samples is two, based on the model DIC and the log-pointwise predictive density (see Table S6). The clustering revealed the potential contribution of each parental species to the nuclear genomic composition of the morphotypes (Fig. 4A). For each morphotype, the average contributions of *K. daigremontiana* and *K. delagoensis* genetic clusters were, respectively: morphotype A (4*x*), 52.97% and 47.03%; morphotype B (3*x*), 35.17% and 64.83%; morphotype C (3*x*), 32.19% and 67.81%; and morphotype D (2*x*), 68.50% and 31.50%.

**FIGURE 4.**
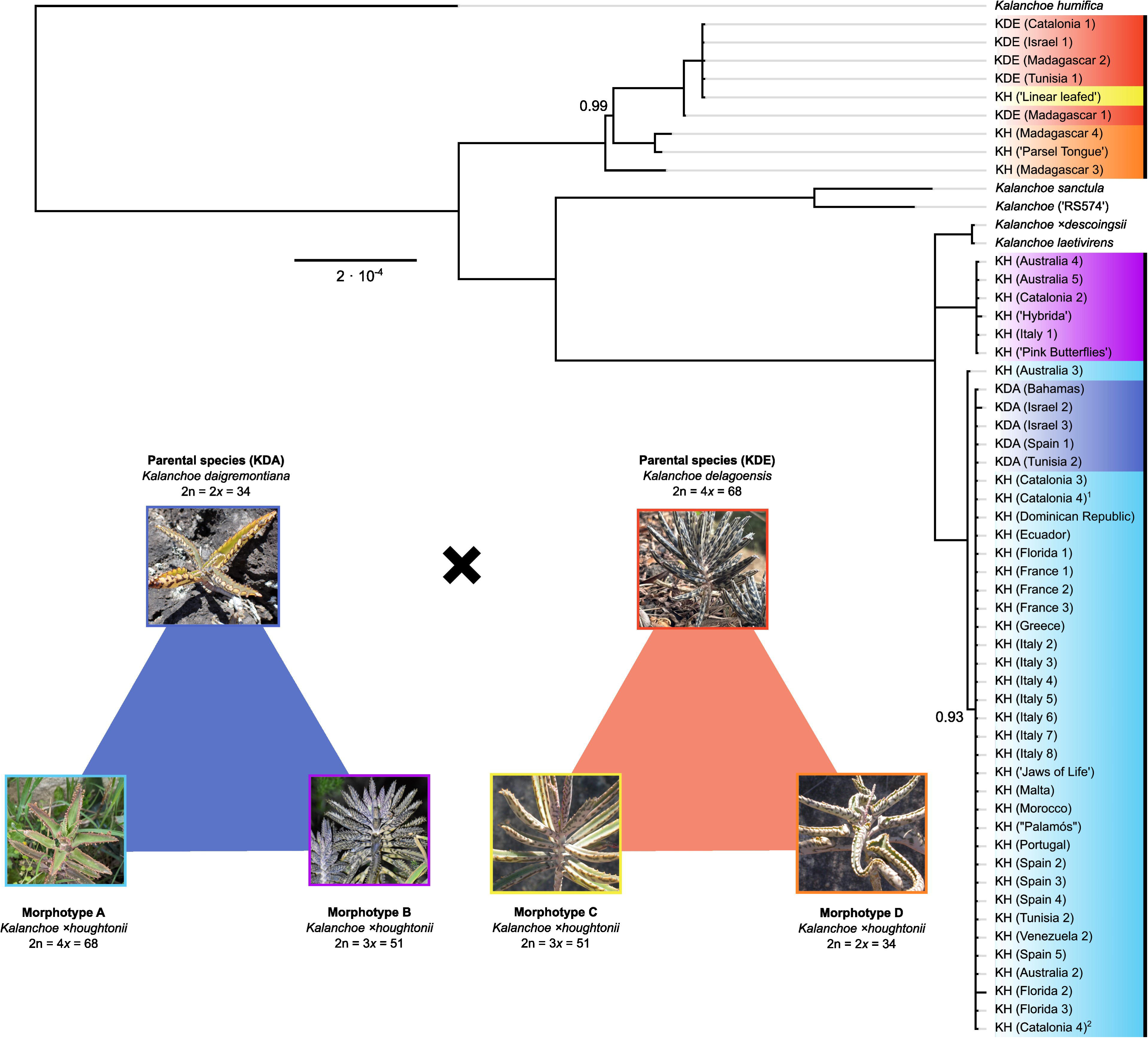
Nuclear genetic variation and structure within the *Kalanchoe ×houghtonii* hybrid complex. (a) Assignment to genetic clusters is shown for K = 2 based on 9,763 bi-allelic SNPs, considering the differential allele dosage in polyploid samples. Technical replicates are indicated with an asterisk. (b) Phylogenetic network based on 1,027 BUSCO genes in 47 *Kalanchoe* samples. (c) A principal component analysis across the first two axes based on 11,966 unlinked bi-allelic SNPs. Parental species and the different *Kalanchoe ×houghtonii* morphotypes are represented in different colours,. Herbarium vouchers are indicated with squared shaped dots.

The nuclear Neighbor-net, constructed using uncorrected P-distances with nuclear genomic data (Fig. 4B), supported the results obtained from the population structure analysis, as well as from the other phylogenomic trees generated in this study (Fig. 3 and Fig. S1). The network clearly separated each hybrid morphotype and the parental populations, linking them through reticulated connections. Among the parental species, there was more genomic variation within *K. delagoensis* than within *K. daigremontiana* samples, as all populations of the latter split from the same node in the phylogenomic network. Regarding the *K.* ×*houghtonii* hybrid complex, the samples of each morphotype were grouped in distinct clusters. In morphotype A, the group with the largest number of samples, the nuclear genomic variation was extremely low, as all samples emerged from the same node, indicating minimal genetic distance between the different populations. However, population “Spain 5” and a replicate of the population “Catalonia 4” from morphotype A, both obtained from herbarium vouchers, are slightly separated from the rest of populations. Morphotype B, although showing reticulations with *K. delagoensis*, clearly constitutes an independent group. Morphotype C was separated from the other three *K.* ×*houghtonii* morphotypes, being as well closely related to *K. delagoensis*. Conversely, all samples from morphotype D appeared more closely related to *K. daigremontiana*.

The principal component analysis (PCA) largely confirmed groupings by morphotype, supporting our previous results. The first and second PC axes respectively explained 23.6% and 18.9% of the variation in the data (Fig. 4C) while adding a third component meant that the first three PC axes accounted for 55.40% of the variation in the data (Fig. S2). In the resulting PCA, the parental populations were respectively distributed in the top-center and the bottom-left sides of the plot, with the different hybrid taxa being gradiently sorted between them. All samples from morphotype A were placed in the top-left quadrant, clearly separated from the rest of *K. ×houghtonii* morphotypes. Samples from morphotypes B and C were more closely related to *K. delagoensis* (bottom left quadrant) while samples from morphotype D were clearly distanced from the rest of morphotypes and parental populations (bottom right quadrant). All samples from each morphotype and parental species were grouped closely, consistently with the results from the population structure and the Neighbor-net analyses. However, some herbarium voucher samples of morphotype A (Florida 1 and Spain 5), morphotype B (Australia 4) and of the parental *K. daigremontiana* (Bahamas) were slightly separated from their core group, likely due to degradation associated with poor DNA preservation. Altogether, these results indicated very low genetic variation within each group, except for *K. delagoensis*, where a sample from Madagascar was clearly separated from the rest, and *K. ×houghtonii* morphotype D, where populations appear well separated.

The assessment of clonality is described in Fig. 5, using the shared heterozygosity (SH) index and the Identity-by-Descent (IBD) analysis. According to the SH index (Fig. 5A), morphotypes A and B showed high signs of clonality (SH index values > 0.90) with the exception of population Florida 1 from morphotype A, for which pairwise comparison values are below the clonality threshold. All pairwise comparisons between populations from morphotype D are below the clonality threshold, showing no evidence of clonality according to SH index. IBD analysis showed similar results (Fig. 5B). Results indicate that all populations of morphotypes A and B would be clonal (IBD values 0.96-0.98 and 0.97-0.99, respectively), excluding Florida 1 from morphotype A (IBD value of 0.94), a sample with significantly degraded DNA signal. Populations of morphotype D have IBD values ranging from 0.79 to 0.85, indicating no clonal origin. Clonality was also detected in both parental species, where all *K. daigremontiana* populations have an IBD value of 0.99, while in *K. delagoensis*, clonality was detected among populations “Israel 1”, “Tunisia 1”, “Madagascar 2” and “Catalonia 1” (IBD values 0.97-0.98), while in population “Madagascar 1” clonality was not detected (IBD values 0.82-0.83).

**FIGURE 5.**
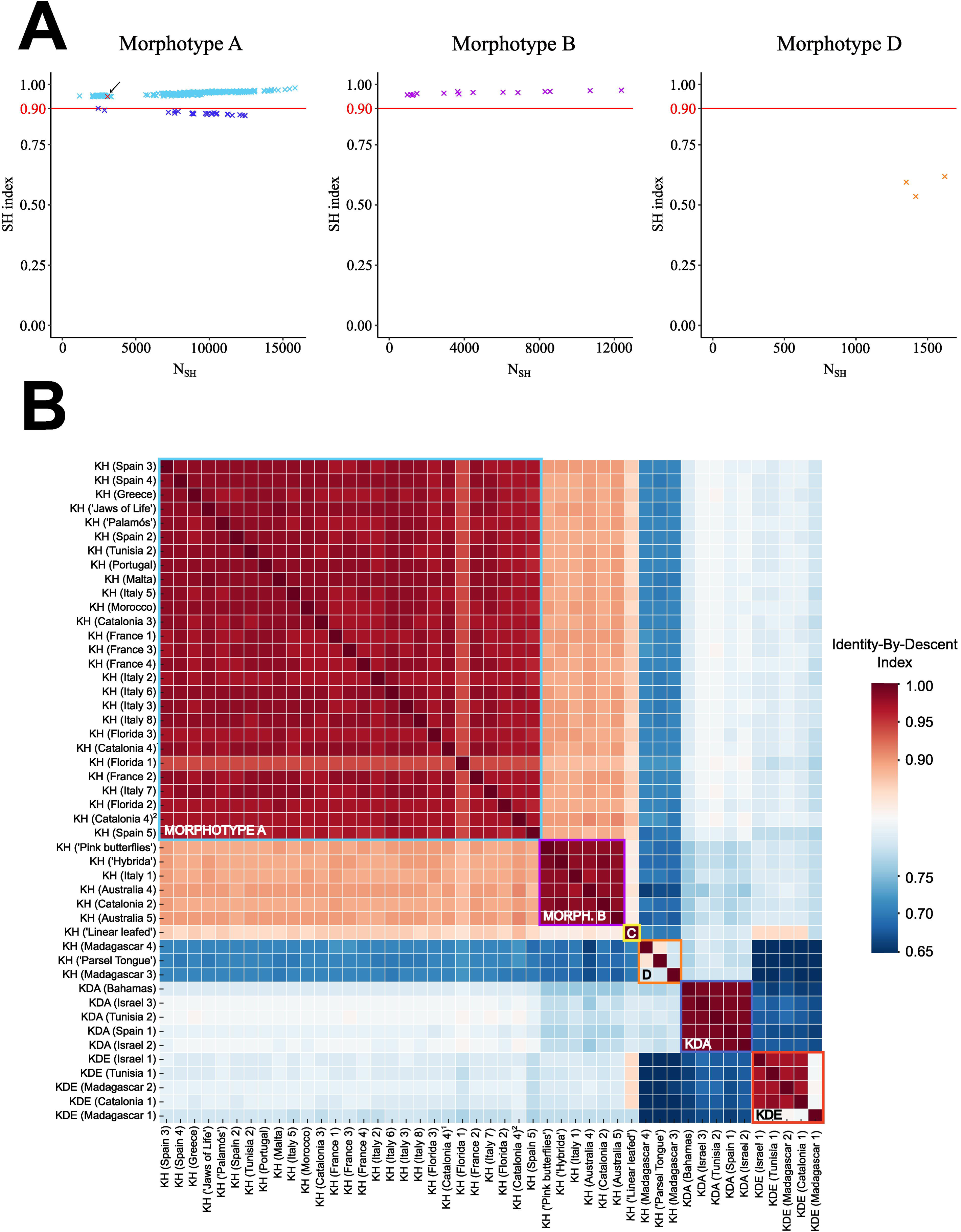
Detecting clonemates pairs using (A) the shared-heterozygosity (SH) index based on 44,055 bi-allelic SNPs. Sample pairs representing technical replicates are marked in red dots and an arrow. In Morphotype A, pairs involving population KH (Florida 1) are indicated in dark-blue. Clonesmates pairs were also detected by (B) the Identity-By-Descent (IBD) index based on 11,966 unlinked bi-allelic SNPs. The empirical IBD cut-off value of 0.95 is defined to consider clonality between pair comparisons.

## Discussion

### Artificial hybridisation in *Kalanchoe ×houghtonii*

According to Shaw *et al*. (2008) and Shtein *et al*. (2021), both morphotypes A (4*x*) and B (3*x*) could have been artificially produced more than once on different continents. Various cultivars have also been described for each of these reportedly artificial morphotypes (Guillot *et al*., 2014; Shtein *et al*., 2021). In theory, the multiple origins and diverse cultivars within these morphotypes could result in substantial genomic and cytogenetic variability of wild populations. However, our findings suggest that the genetic variability present in populations corresponding to the reportedly artificial morphotypes A and B is surprisingly low. Specifically, plastome sequences (Fig. 3) and nuclear genomic data (Fig.s 4 and Fig. S1) indicate that all analysed populations across the Mediterranean Basin, Australia and America from morphotype A—tetraploid, as hypothesised by Shtein *et al*. (2021) and confirmed cytogenetically in this study for the first time (Table 2)— share the same genotype, confirming its presence in the wild at least since 1966 (Australia). To validate these results, we included two technical replicates of population “Catalonia 4” of morphotype A in our analyses, with the results suggesting that the slight genetic differences observed between populations of morphotype A are probably non-significant and can be attributed to sequencing errors and/or somatic mutations. Our findings are consistent, on a global scale, with those obtained by Guerra-Garcia *et al*. (2015) in a population genetics study on four invasive Mexican populations of *K. ×houghtonii*. Employing microsatellite markers, these authors found that a single genotype of the hybrid had been successfully introduced and further expanded in the region by clonal growth. In our current study, using genome-level markers and a global sampling, we show that a single clone of *K. ×houghtonii* has been able to colonise and invade mild-climate regions in four continents.

Genetic variation and evolution are expected to play an important role in the success of invasive species (Dlugosch and Parker, 2007). Despite the hypothesis that invasive species may experience decreased genetic variation due to population bottlenecks during colonisation events (Hollingsworth and Bailey 2000; Allendorf and Lundquist 2003: Poulin *et al*., 2005; Wang *et al*., 2005; Roman and Darling, 2007; Li *et al*., 2006; Lambertini *et al*., 2010), some IAPS populations harbour high genetic diversity (Pappert *et al*., 2000; Meekins *et al*., 2001; Maron *et al*., 2004; Genton *et al*., 2005; Gutierrez-Ozuno *et al*., 2009). However, in *K.* ×*houghtonii*, the widespread and aggressive invader morphotype A of our study lacks genomic variability and just one clonal genotype has been observed in all sampled populations of this morphotype (Fig. 4, Fig. 5, Fig. S1). These findings match the hypothesis of the “general-purpose-genotype” (Baker 1955, 1974; Ferrero *et al*., 2015), which suggests that the most successful coloniser would be the one to thrive without genetic variation, relying instead on a single “best genotype” capable of colonising a wide variety of environments (Dlugosch *et al*., 2016). Shtein *et al*., (2021) reported that the apparent morphological differences between the cultivars ‘Jaws of Life’ and ‘J.T. Baldwin’ of morphotype A disappeared when grown under the same conditions. In our genomic study, we have observed that samples named ‘Garbí’ and ‘J.T. Baldwin’ of morphotype A (i.e. those wild-growing in Europe or in America, respectively) correspond to the same genotype. Therefore, the reported morphological differences among these cultivars are likely due to the wide phenotypic plasticity of this invasive genotype.

Although our results prove that morphotype A has a single origin, its exact source remains unclear. Initially, it had been proposed that morphotype A could have been created by Houghton (1935) or by Resende (1954). Our genomic data shed light on this question, with all genotyped samples of morphotype A displaying the same plastome haplotype found in all analysed *K. daigremontiana* populations, revealing that this species was the maternal species in the cross. Hougthon (1935) also reported that the maternal species in the cross between the parental species was *K. daigremontiana*. Yet, clones originating from Houghton’s crossings, described as “identical plants”, corresponded to morphotype B. Moreover, samples from Houghton’s gardens in San Fernando (California) were later determined cytogenetically as triploids by Baldwin (1949). Resende (1954) reported a single tetraploid clone resulting from the cross between *K. delagoensis* and *K. daigremontiana*, with *K. delagoensis* as the maternal genome donor. Therefore, our results reject the hypothesis that the origin of morphotype A is either the cross reported by Houghton (1935) or by Resende (1954). The actual source of this morphotype is still a mystery, and may be either of artificial (as a result of some crossbreeding that has not been reported) or of natural origin (arising from spontaneous crossings in native or non-native areas where both parentals coexist; see Ward 2006; Herrando-Moraira *et al*., 2020), similarly as it occurred in Madagascar with morphotypes C and D (see below). The fact that there has not been any single report, up to our knowledge, of morphotype A plants in Madagascar (neither registered in local herbaria, nor in GBIF or iNaturalist), in spite of both parental species coexisting in certain areas, argues against the possibility of a natural origin. Besides, crosses between plants of different ploidy levels would often result in failure of endosperm development (Birchler, 2014). Morphotypes B and C are triploid (2*n* = 3*x* = 51), the most likely expected outcome from a cross between a diploid and a tetraploid. The cytogenetic composition of morphotype A, a tetraploid hybrid (2*n* = 4*x* = 68), is even more exceptional, as it implies necessarily the presence of a non-disjunct gamete from the diploid genome donor: the resulting hybrid has *n* = 34 from *K. daigremontiana* (2*n* = 2*x* = 34, *n* = 34 gametes) and *n* = 34 from *K. delagoensis* (2*n* = 4*x* = 68, *n* = 34 gametes). However, although artificial crossing could have made it easier to produce and select tetraploid plants, natural hybridization in localities where both parentals coexist cannot be discarded.

### Origin of natural hybrid forms in *Kalanchoe ×houghtonii*

Plants representing morphotype C (3*x*) are difficult to distinguish from *K. delagoensis*, and they occur naturally in Madagascar, for instance, along the Onilahy River, where the ranges of *K. daigremontiana* and *K. delagoensis* overlap (Shtein *et al*., 2021). Based on morphological and distribution data, plants of this morphotype have been proposed either to be a closely related non-hybrid taxon showing intermediate characters between the two parental species or to represent a natural hybrid. Our genomic population analysis indicates that plants of morphotype C show the genomic composition expected for a triploid cytotype derived from the cross between these parental species (i.e. one third of the genome representing the diploid *K. daigremontiana* and two thirds representing the tetraploid *K. delagoensis*) (Fig. 4A). In morphotype C, the cross between the parental species is reversed as compared to plants representing morphotypes A and B, where *K. delagoensis* is the maternal genome donor and *K. daigremontiana* the paternal donor. The direction in which a hybrid cross is made has strong consequences for the transcriptional program of offsprings (Joseph *et al*., 2015; Flood *et al*., 2020). Maternal effects are based on physiological properties expressed in the mother plant, which are passed on to their progeny (Botet and Keurentjes, 2020). Hence, this switch in the crossing between both parentals could explain why plants representing morphotype C are rather morphologically similar to *K. delagoensis* as compared with plants of morphotype A and B, for which the maternal donor is *K. daigremontiana*.

The last described variant of *K. ×houghtonii* is morphotype D, for which we report the diploid ploidy level for the first time (Table 2). According to Shtein *et al*. (2021), plants referable to *K. ×houghtonii* morphotype D are very variable, from almost indistinguishable from *K. daigremontiana* to plants clearly showing intermediate characters between *K. daigremontiana* and *K. delagoensis*. These plants occur naturally in Madagascar, or in cultivation (e.g. cultivar ‘Parsel Tongue’). Our results suggest that the morphological variability found in plants representative of the morphotype D could be related to their genetic and genomic diversity. We detected two different plastome sequence haplotypes corresponding to the parental species *K. delagoensis*, being this taxon the maternal donor of this morphotype. The genetic diversity within the nuclear genome also shows significant differentiation among samples and confirms that *K. daigremontiana* is the highest contributor in the genome composition of this morphotype (Fig. 4, Fig. 5). Besides, clonality assessments (Fig. 5) confirm that, unlike the other morphotypes, morphotype D populations would not have clonal origin. Hence, both plastome and nuclear data support the hybrid origin of this morphotype. Because of the morphological similarity between *K. daigramontiana* and morphotype D samples, it has been suggested that this morphotype—specifically, the cultivar ‘Parsel Tongue’— may represent a back-cross of *K. daigremontiana* with a fertile cultivar of *K. ×houghtonii* ‘J.T. Baldwin’ (Shtein *et al*., 2021). However, considering that plants of morphotype A (including the cultivar ‘J.T. Baldwin’) are tetraploid, and *K. daigremontiana* plants are diploid, the direct combination of gametes from the tetraploid and the diploid would lead to triploid or tetraploid progeny. Furthermore, the inheritance of the plastome sequence must be considered, since morphotype A samples have *K. daigremontiana* as maternal donor of chloroplast genome, while plants representing morphotype D have *K. delagoensis* plastome instead. Therefore, the hypothesis of morphotype D originating as a back-cross of *K. daigremontiana* with a fertile cultivar of *K. ×houghtonii* morphotype A can be rejected because of its diploid ploidy level and the inheritance of the chloroplast genome. According to our results, introgression between *K. daigremontiana* (2*x*) and an unknown hybrid taxon containing *K. delagoensis*-like plastome—such as morphotype C, which shows connections with morphotype D in nuclear phylogenomic networks, and coexists with it in Madagascar— might be a plausible origin.

### Invasiveness of the *Kalanchoe ×houghtonii* hybrid complex and beyond

*Kalanchoe ×houghtonii* is an exemplary biological system to study the effects of hybridisation and polyploidy on the emergence of invasive taxa. The hybridogenic nature of this complex is clearly shown, with four hybrid taxa from different genomic combinations from the same parents, different ploidy levels and showing completely different cytogenomic features and invasive abilities. Hybridisation is common in *Kalanchoe* (Kuligowska *et al*., 2015), involving in many cases *K. delagoensis* as a parental species (Shtein *et al*., 2021). For instance, *K. ×descoingsii*, a recent hybrid taxon found in Antsokay Arboretum of Madagascar has been suggested (based on plant morphology) to be a cross between *K. delagoensis* and *K. laetivirens* (Shtein *et al*., 2021). This is evident in the ribosomal DNA networks (Fig. S1), where the 5S rDNA corresponds to that of *K. laetivirens* while the 35S rDNA is closely related to *K. delagoensis*. Besides, our results derived from the plastome sequence analyses suggest that *K. laetivirens* is the maternal genome donor species of *K. ×descoingsii* (Fig. 1). This hybrid can reach its bulbil producing phase rapidly and it spreads easily, similarly to *×houghtonii* (Shtein *et al*., 2021). Based on ploidy level estimations provided in this study (Table 2), *K. ×descoingsii* might be a pentaploid, while *K. laetivirens* is potentially a hexaploid, the results being consistent with the expected crossing between *K. delagoensis* (4*x*) and *K. laetivirens* (6x).

In the context of global change, with rapidly changing environments and increasingly frequent extreme climatic events, generalist species with high phenotypic plasticity hold a significant biological advantage over specialist species (Clavel *et al*., 2010). As already proposed in *Spartina* (Ainouche *et al*., 2009), our study shows that hybridisation and polyploidy can be important sources of generalist genotypes, which may eventually displace more specialist taxa. The interaction between clonal growth and climate change has also begun to attract interest (Yu *et al*., 2016), being the most problematic IAPS in this context those showing clonal propagation (Pysek, 1997; Cadotte *et al*., 2006; Roiloa 2019). Clonal reproduction, polyploid genomes and absence of genetic diversity seems to be a recipe for successful invasiveness.

Altogether, the *K. ×houghtonii* complex exemplifies a model for understanding hybrid plant invasions, especially in the context of global change marked by drought, erosion, and rising temperatures (Seneviratne *et al*., 2021). Its multiple cytotypes with varying colonizing capacities, high morphological plasticity, and resilience to drought and heat contribute to its invasiveness. Its high rates of clonal reproduction, but in particular, the presence of a single “general purpose genotype” responsible for its global spread, add to a suite of traits enhancing its invasiveness. The case of the *K. ×houghtonii* hybrid complex, whose origin is linked to horticultural practice, advocates for a “climate-smart invasive species management” as suggested by Colberg *et al*. (2024). Mitigating the hybrid and parentals spread by limiting their sale and transport, adopting preventive actions, as well as establishing a regulatory framework across those countries environmentally more sensitive to their invasion (mostly, but not limited to those of Mediterranean climate) can be crucial for addressing the spread of this complex which has reached all continents (except Antarctica) in less than a century.

## Acknowledgements

The following herbaria are acknowledged for their support in providing several *Kalanchoe* vouchers: Queensland Herbarium (Australia), University of Florida Herbarium, The New York Botanical Garden, Missouri Botanical Garden Herbarium (USA), and the BC herbarium (Spain). The Supercomputing Center of Galicia (CESGA) supplied the necessary computation resources for data processing. Finally, we want to express our gratitude to the following people who helped in different aspects of this research: Núria Abellán, Víctor Álvarez, Manica Balant, Lucas C. Majure, Teresa Garnatje, Neus Ibáñez, Carlos Gómez-Bellver, Eduard López-Guillén, Vanessa Lozano, Joan Ramon Mendo, Stephen Mifsud, Jaume Pellicer, Jaume Pàmies, Lisa Pokorny, Samuel Pyke, Gideon Smith, Jaume X. Soler, Eva M. Temsch and Miquel Veny.

## Author contributions

J.L-P, S.G. and D.V. conceived the study. J.P.P-D, J.L-P, N.N., I.P.L, R.S. and S.G collected samples. J.P.P-D, N.B. and L.V. carried out laboratory work. J.P.P-D, N.B., S.G. and D.V. analysed data. J.P.P-D, S.G., and D.V. wrote the first version of the manuscript. All authors contributed in interpreting results and edited and approved the final version of the manuscript.

## Data availability statement

The data that support the findings of this study are openly available in NCBI at https://www.ncbi.nlm.nih.gov/bioproject/PRJNA944709/ reference number PRJNA944709. All ribosomal RNA genes and the entire plastome sequences are openly available in the NCBI repository GenBank (see Table S1).

---

**FIGURE S1.**
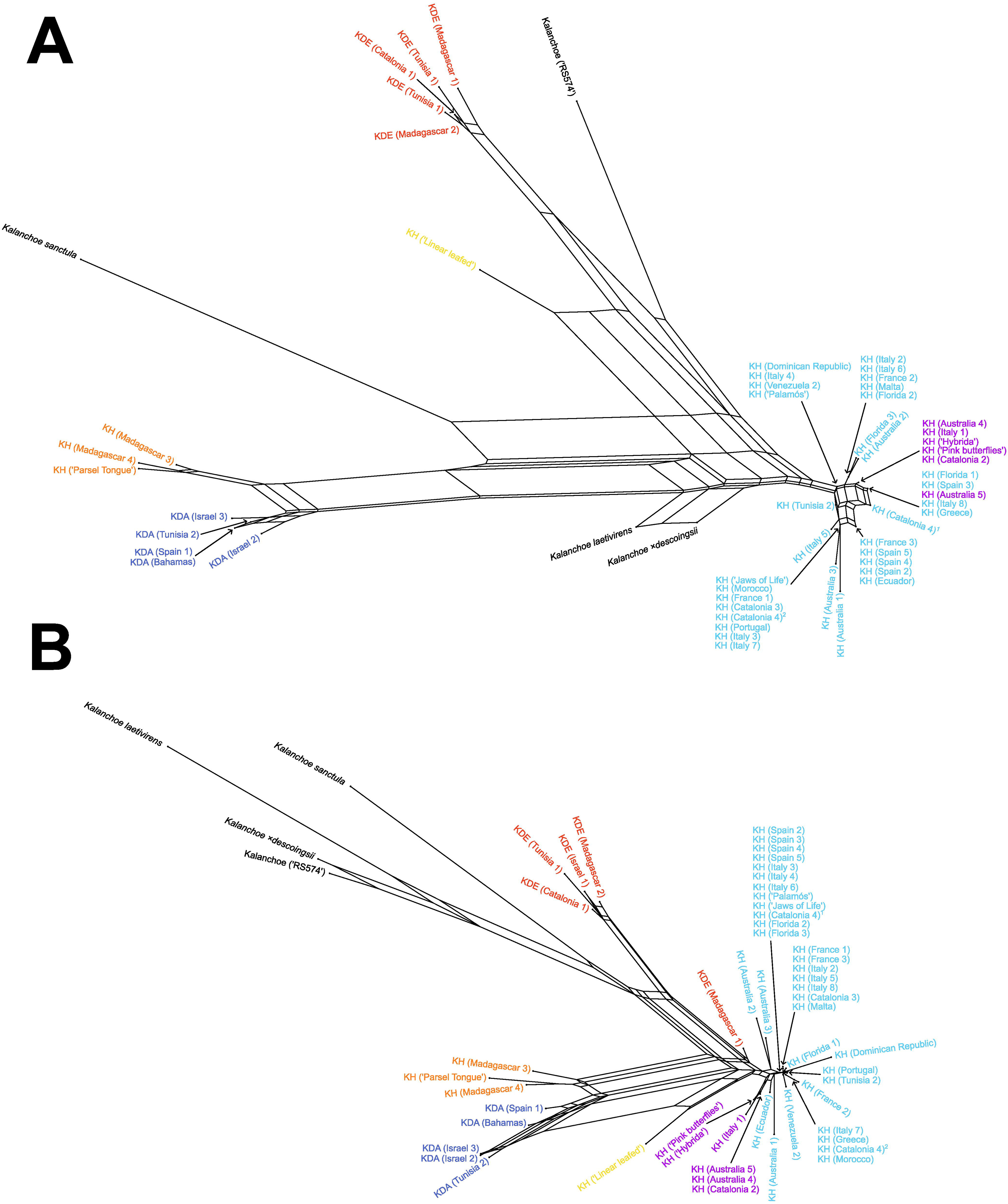
Evolutionary relationships based on rDNA among the different *Kalanchoe ×houghtonii* morphotypes, its parentals and other closely related species and hybrid complexes. Neighbor-Net inferred from (a) the 5S gene and intergenic spacer and (b) the ITS1, ITS2 intergenic spacers and 35S ribosomal DNA genes.

**FIGURE S2.**
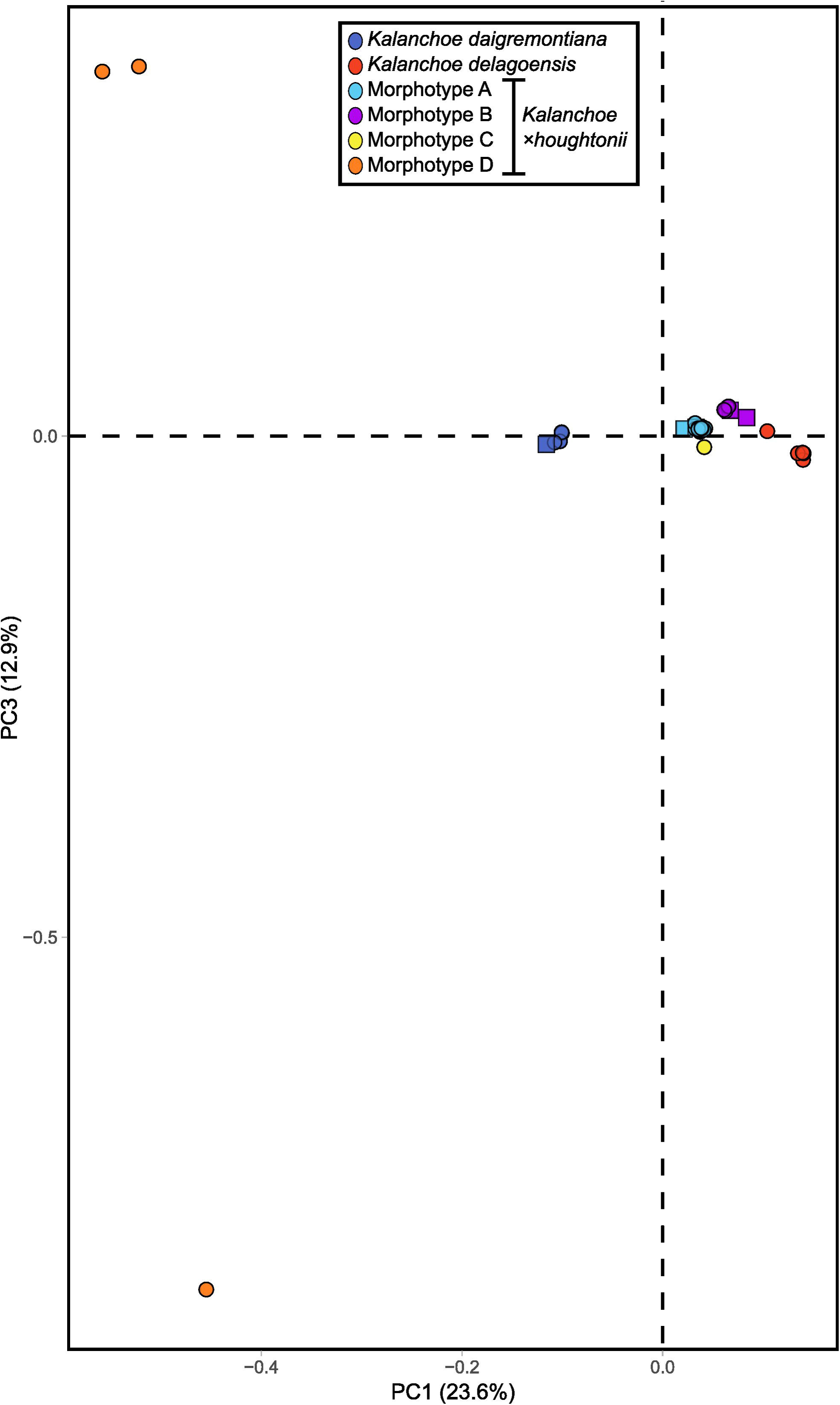
Principal component analysis across the first and the third axes, with the genetic group coloured based on the *Kalanchoe* taxa.

**FIGURE S3.**
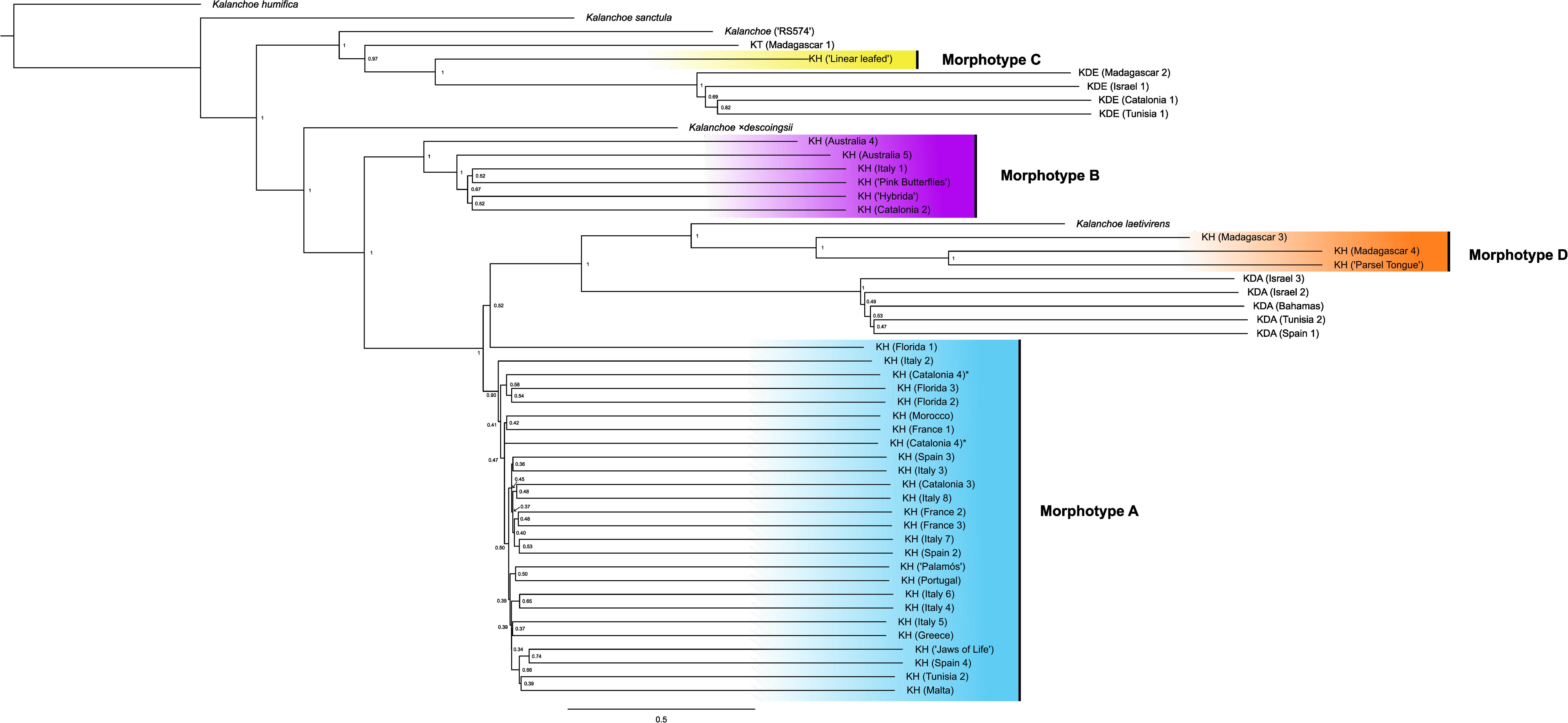
Light microscope photographs of metaphase chromosomes of: (a) *K. daigremontiana* (2n=4x=*ca.*34); (b) *K.* ×*houghtonii* (triploid, 2n=3x=*ca.* 51); (c) *K.* ×*houghtonii* (tetraploid, 2n=4x=*ca.* 68); (d) *Kalanchoe tubiflora* (2n=4x=*ca.* 68)

**FIGURE S4.**
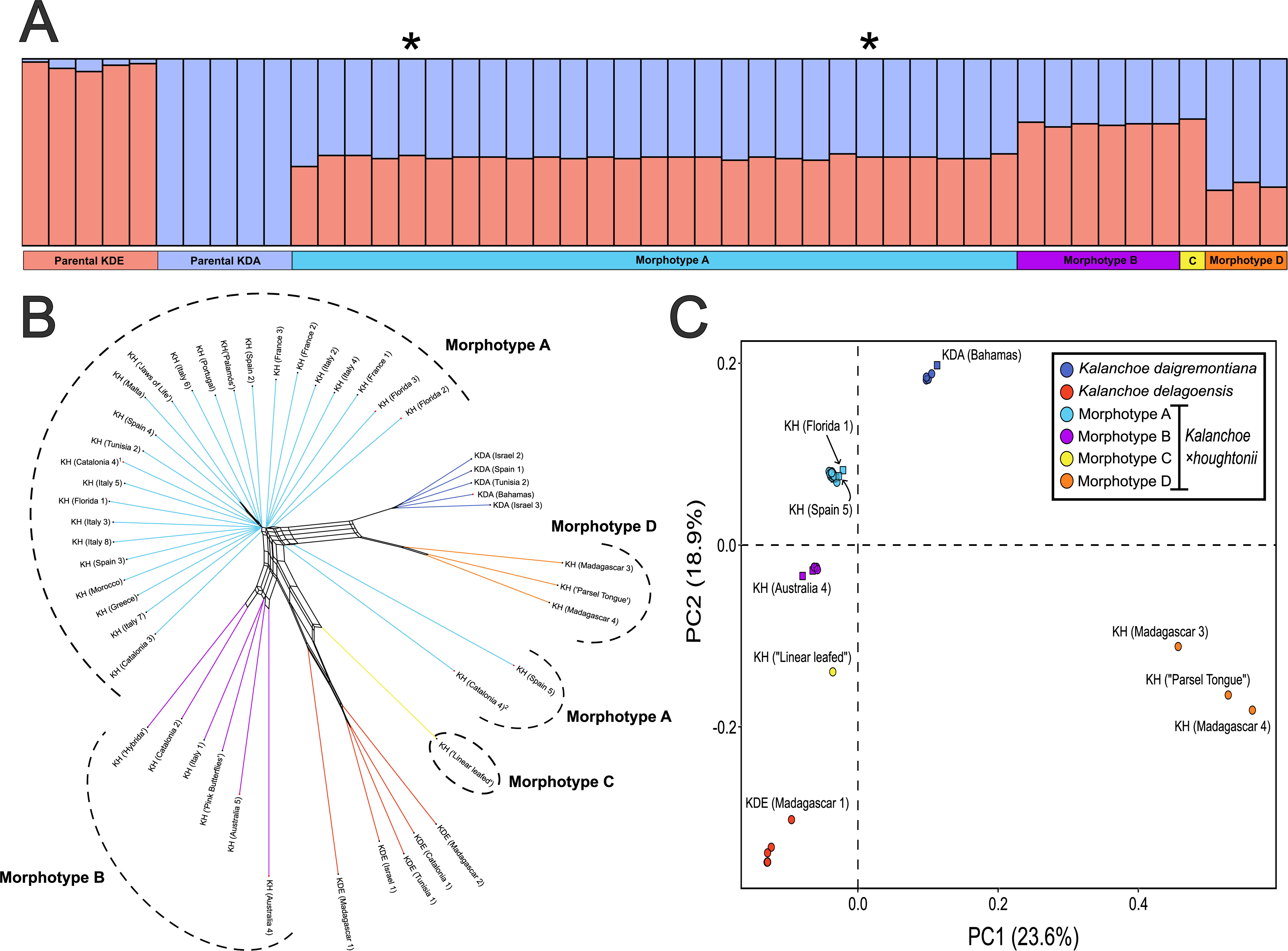
Phylogenomic tree of 50 samples of *Kalanchoe* based on 1,122 filtered BUSCO genes, using as outgroup the species *Kalanchoe humifica*. The tree includes all morphotypes of *Kalanchoe ×houghtonii* (A, B, C and D), the parental species (*K. daigremontiana* and *K. delagoensis*), and other related taxa (*K. sanctula*) and hybrid complexes (*K. laetivirens* and *K. xdescoingsii*). Samples of *Kalanchoe* ×*houghtonii* morphotype A are indicated in light blue, morphotype B in purple, morphotype C in yellow and morphotype D in orange. Samples of parental species are indicated in dark blue for *Kalanchoe daigremontiana* and magenta for *Kalanchoe delagoensis*. Nodes without support values are considered maximum supported (pp = 1).

**TABLE S1.** Information of all *Kalanchoe* species and populations sampled in this study, including the collectors, the herbarium vouchers, the SRA codes and the GenBank accessions of the plastome and the ribosomal DNA sequences.

**TABLE S2.** Descriptive statistics of the number of recovered genes, guanine-cytosine percentage, mapping details and GenBank accessions of all plastome reconstructions of each *Kalanchoe* population sampled.

**TABLE S3.** Detailed table summarising target gene recovery efficiency from HybPiper analyses. Table includes information about the number of input reads mapped to target genes, the number of genes in the target file that had reads mapped, and the number of genes with sequences >25, 50 and 75% of the mean target length.

**TABLE S4.** Detailed table summarising the statistics from *max_overlap* pipeline of capture coverage of target sequences rescued by HybPiper. Information of representedness (proportion of species for which genes were obtained), completeness (proportion of gene sequences obtained for each species) and evenness (how evenly the sequence lengths are distributed across species) is provided.

**TABLE S5.** Detailed alignment statistics provided by AMAS for the preliminary alignment and the subsequent corrected alignment using TreeShrink with two different false positive tolerance levels (0.01 and 0.05).

**TABLE S6.** Information about the deviance information criterion (DIC) and the log-pointwise predictive density for each one of the clusters (K) tested with the ENTROPY analyses.

